# When is a control not a control? Reactive microglia occur throughout the control contralateral visual pathway in experimental glaucoma

**DOI:** 10.1101/853275

**Authors:** James R Tribble, Eirini Kokkali, Amin Otmani, Flavia Plastino, Emma Lardner, Rupali Vohra, Miriam Kolko, Helder André, James E Morgan, Pete A Williams

**Affiliations:** Department of Clinical Neuroscience, Division of Eye and Vision, St. Erik Eye Hospital, Karolinska Institutet, Stockholm, Sweden; School of Optometry and Vision Sciences, Cardiff University, Cardiff, Wales, U.K.; Department of Veterinary and Animal Sciences, Pathobiological Sciences, University of Copenhagen, Denmark; Department of Drug Design and Pharmacology, University of Copenhagen, Copenhagen, Denmark; Department of Ophthalmology, Rigshospitalet-Glostrup, Copenhagen, Denmark; School of Medicine, Cardiff University, Cardiff, Wales, U.K.

**Author notes:** Correspondence to: Pete A Williams, Department of Clinical Neuroscience, Division of Eye and Vision, St. Erik Eye Hospital, Karolinska Institutet, Stockholm, Sweden; @pete_the_teapot. Authors contributed equally to this work (James R Tribble, Eirini Kokkali). (JRT), (EK), (AO), (FP), (EL), (RV), (MK), (HA), (JEM), (PAW).

**Keywords:** microglia, monocyte, neuroinflammation, glaucoma, retina, optic nerve, head, lateral geniculate nucleus, superior colliculus

## Abstract

**Purpose:** Animal models show retinal ganglion cell injuries that replicate features of glaucoma and the contralateral eye is commonly used as an internal control. There is significant cross-over of retinal ganglion cell axons from the ipsilateral to the contralateral side at the level of the optic chiasm which may confound findings when damage is restricted to one eye. The effect of unilateral glaucoma on neuroinflammatory damage to the contralateral visual pathway has largely been unexplored.

**Methods:** Ocular hypertensive glaucoma was induced unilaterally or bilaterally in the rat and retinal ganglion cell neurodegenerative events were assessed. Neuroinflammation was quantified in the retina, optic nerve head, optic nerve, lateral geniculate nucleus, and superior colliculus by high resolution imaging, and in the retina by flow cytometry and protein arrays.

**Results:** Following ocular hypertensive stress, peripheral monocytes enter the retina, and microglia become reactive. This effect is more marked in animals with bilateral ocular hypertensive glaucoma. In rats where glaucoma was induced unilaterally there was significant microglia activation in the contralateral (control) eye. Microglial activation extended into the optic nerve and terminal visual thalami, where it was similar across hemispheres irrespective of whether ocular hypertension was unilateral or bilateral.

**Conclusions:** These data suggest that caution is warranted when using the contralateral eye as control in unilateral models of glaucoma.

**Translational Relevance:** Use of a contralateral eye as a control may confound discovery of human relevant mechanism and treatments in animal models. We also identify neuroinflammatory protein responses that warrant further investigation as potential disease modifiable targets.

## Introduction

Glaucoma is the leading cause of irreversible blindness, affecting ∼80 million people worldwide (1). Glaucoma is characterized by the progressive dysfunction and loss of retinal ganglion cells and their axons. Neuroinflammation is a shared feature of glaucoma pathogenesis in human glaucoma patients (2) and animal models of glaucoma (genetic and inducible (3)). In times of stress, retinal ganglion cells are reliant on a supportive network of glial cells to provide neurotrophic and metabolic support, especially to their long, unmyelinated axons in the retina and optic nerve head. Emerging evidence supports a paradigm in which retinal ganglion cell axons are insulted throughout their trajectory to retinorecipient areas; the dorsal lateral geniculate nucleus and the superior colliculus. Microglia and/or other immune-derived cells are likely to be key players in these pathogenic events.

The eye is commonly used as a tool to explore neurodegenerative events. However, a common feature of these experiments is the use of the contralateral eye as an internal control (which is often used to normalize data to the experimental eye). These data should be qualified because: *i*) retinal ganglion cell numbers differ between eyes even within individual animals (4), *ii*) significant numbers of retinal ganglion cell axons do not cross at the optic chiasm and terminate in ipsilateral thalami (ranging from ∼3% in mouse and ∼3-10% in rat to ∼45% in primates and ∼50% in human (5-7)), *iii*) albino rodents are commonly used (*e.g.* Wistar and Sprague-Dawley rats) which have reduced ipsilateral projections (8-11), and, *iv*) most human primary open angle glaucoma is bilateral. Microglial activation is an early and persistent feature across glaucoma models and species with evidence of immune-cell activation in human tissue from advanced glaucoma (12-16). To date, there is a paucity of data regarding microglial activation in the contralateral eye of mice with unilateral ocular hypertension (largely qualitative with quantification only through cell counting (17-19)). Activation has been reported in the optic nerve and in terminal brain thalami (14, 20, 21), but contralateral activation has yet to be fully explored in these tissues.

To address these issues and explore any neuroinflammation throughout the visual pathway in glaucoma, we fully characterize inflammatory changes in an inducible rat model of ocular hypertensive glaucoma in which intraocular pressure is elevated by intracameral (anterior chamber) injection of paramagnetic beads to occlude the drainage structures of the eye (22, 23). This results in sustained ocular hypertension driving a reproducible and significant loss of retinal ganglion cell numbers as well as neuroinflammation (without uveitis). Using this model, we assessed the effects of unilateral or bilateral induced ocular hypertension on glaucoma pathology throughout the retinal ganglion cell projections. These experiments were performed in the Brown Norway rat as it is the genetic standard for rats, is inbred, pigmented, and avoids retinothalamic pathway issues present in albino animals. We demonstrate that neuroinflammation persists throughout retinal ganglion cell pathways with monocyte infiltration and pro-inflammatory cytokine release in the retina. Critically, these changes are mirrored in the contralateral visual pathway warranting caution when using the contralateral eye as a ‘control’ in ophthalmic research.

## Methods

### Rat strain and husbandry

Adult, male Brown Norway rats (aged 12-16 weeks, weighing 300-375 g, SCANBUR) were housed and fed in a 12 h light / 12 h dark cycle with food and water available *ad libitum*. All experimental procedures were undertaken in accordance with the Association for Research for Vision and Ophthalmology Statement for the Use of Animals in Ophthalmic and Research. Individual study protocols were approved by Stockholm’s Committee for Ethical Animal Research (10389-2018).

### Induction of ocular hypertension

Ocular hypertension (OHT) was induced bilaterally (OHT/OHT; *n* = 22 rats, 44 eyes) or unilaterally (OHT/NT; *n* = 12 rats, 12 eyes). For unilateral OHT rats, the contralateral eye was an un-operated, normotensive (NT) eye (NT/OHT; *n* = 12 rats, 12 eyes). Bilateral NT rats served as controls (NT/NT; *n* = 17 rats, 34 eyes; **Figure 1A**). OHT was induced using a magnetic microbead injection model as described previously. Microbeads (Dynabead Epoxy M-450, Thermo Fisher) were prepared in HBSS and 6-8 µl injected into the anterior chamber. Beads were distributed with a magnet to block the iridocorneal angle. Intraocular pressure (IOP) was measured using a Tonolab rebound tonometer (Icare) in awake, unrestrained rats, habituated to the tonometry procedure. Baseline IOP was recorded on the day of surgery (day 0) and recorded every 2-4 days afterwards until the endpoint (postoperative day 14), with IOP recordings always taken between 9 and 10am to avoid the effects of the circadian rhythm on IOP (2-3 hours from lights on). IOP was taken as the average of 5 tonometer readings. NT/NT rats followed the same 14 day time course (without surgery).

**Figure 1:**
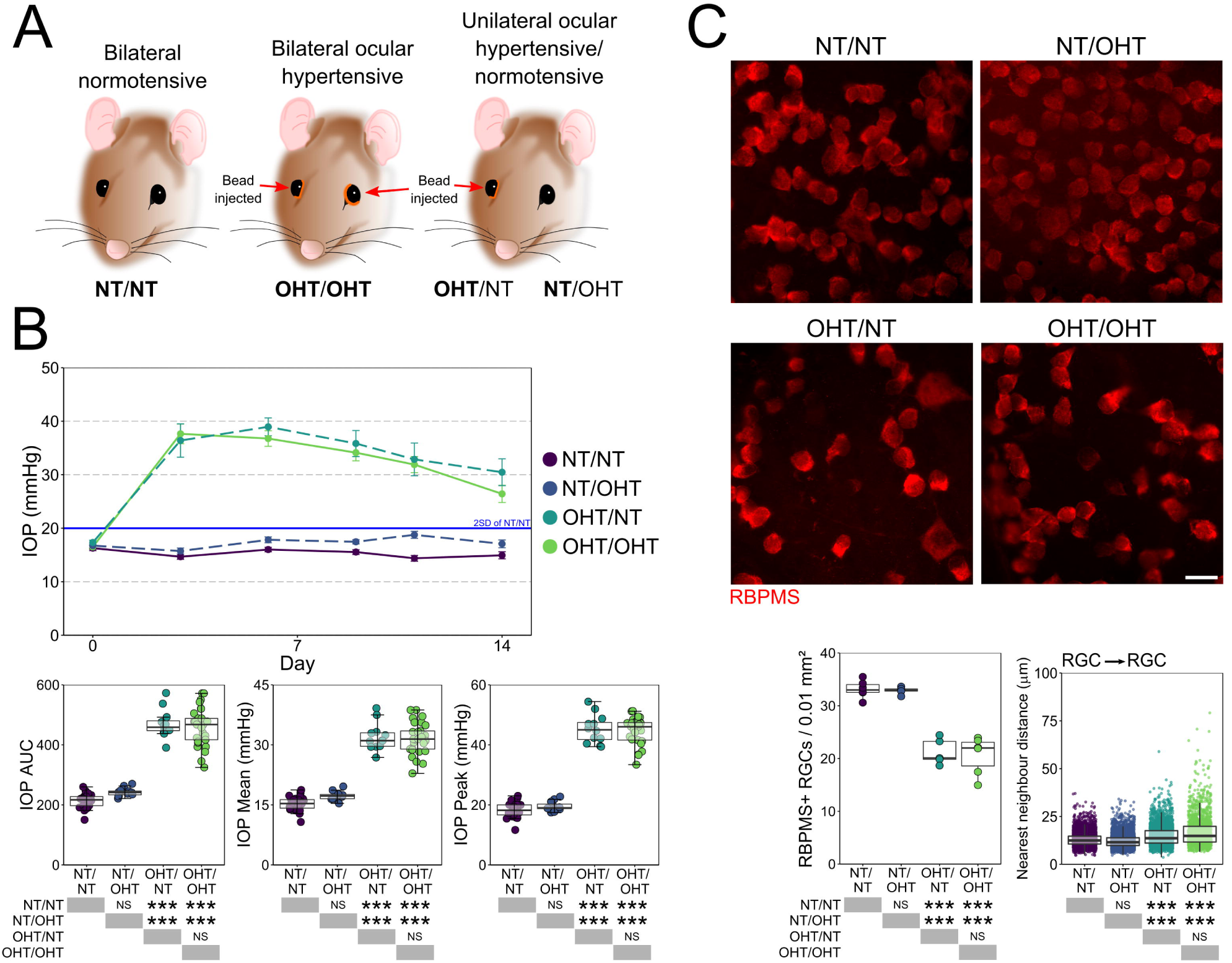
Induction of OHT via paramagnetic bead induction. (**A**) Brown Norway rats underwent intracameral injections of paramagnetic beads either bilaterally or unilaterally. The contralateral eye remained un-operated in unilaterally injected rats. (**B**) Beads were pulled into the drainage structures of the eye via a rare earth magnet resulting in significant and robust IOP increase that was sustained until euthanasia (**B**). 14 days of OHT results in significant RGC atrophy demonstrated in flat-mount retinas confirming a significant loss of RGCs and increased nearest neighbor distance of RGCs following 14 days OHT. For complete *n* please see **Methods**. Scale bars = 20 μm in **C**. * = *P* < 0.05, ** = *P* < 0.01, *** *P* < 0.001, NS = non-significant (*P* > 0.05).

### General histopathology

Rats were heavily anaesthetized at day 14 by intraperitoneal injection of pentobarbital (75 mg/kg), and euthanized by cervical dislocation. Eyes were immediately enucleated (with 1 mm of optic nerve post-globe preserved) and immersed in 3.7% PFA in 1x PBS. Brains (with attached optic nerves) were removed and immersed in 3.7% PFA in 1x PBS. Eyes, optic nerves, and brains for cryo-sectioning were maintained in fixative for 24 hours before cryo-protecting in 30% sucrose in 1x PBS for 24 hours, freezing in optimal cutting temperature medium (Sakura) on dry ice, and stored at -80 °C. Cryo-sections were cut on a cryostat (Cryostar NX70, Thermo Scientific) and stored at -20 °C. Eyes were sectioned at 20 μm thickness (anterior to dorsal plane), with sections maintained from ∼200 μm nasal to ∼200 μm temporal of the optic nerve head (*n* = 3 eyes for NT/NT, OHT/OHT, NT/OHT, and OHT/NT). Optic nerves for cryo-sectioning were maintained as pairs attached at the optic chiasm, and sectioned at 20 μm to generate longitudinal nerve sections (*n* = 4 optic nerves for NT/NT, OHT/OHT, NT/OHT, and OHT/NT). Brains were serial sectioned (coronal plane) at 50 μm from -4 mm caudal of cranial landmark Bregma, to -8 mm caudal of Bregma in order to capture the entire dLGN and SC (*n* = 3 brains [6 independent hemispheres] for NT/NT, OHT/OHT, NT/OHT, and OHT/NT).

### Immunohistochemistry and immunofluorescence

Antibodies and stains used for immunofluorescence, immunohistochemistry, and histological labelling are detailed in **Table 1**. The same immunofluorescent labelling protocol was followed for cryo-sections and flat mount retina with exceptions noted where relevant. Tissue was isolated using a hydrophobic barrier pen (VWR), permeabilized in 0.5% Triton X-100 in 1x PBS for 1 h, blocked in 5% bovine serum albumin in 1x PBS for 1 h, and primary antibody applied overnight at 4 °C (see **Table 1**). Tissue was then washed for 5 × 5 min in 1x PBS, secondary antibody applied (1:500 in 1x PBS) for 4 h at room temperature, and washed for 5 × 5 min in 1x PBS. DAPI nuclear stain (500 μg/ml stock; used 1:500 in 1x PBS) was applied for 10 min, the tissue washed for 5 min in 1x PBS, dried and mounted using Fluoromount-G and glass coverslips (Invitrogen). For cryo-sections, tissue was air dried for 15 min and rehydrated in 1x PBS for 15 min before following the protocol above.

**Table 1.**
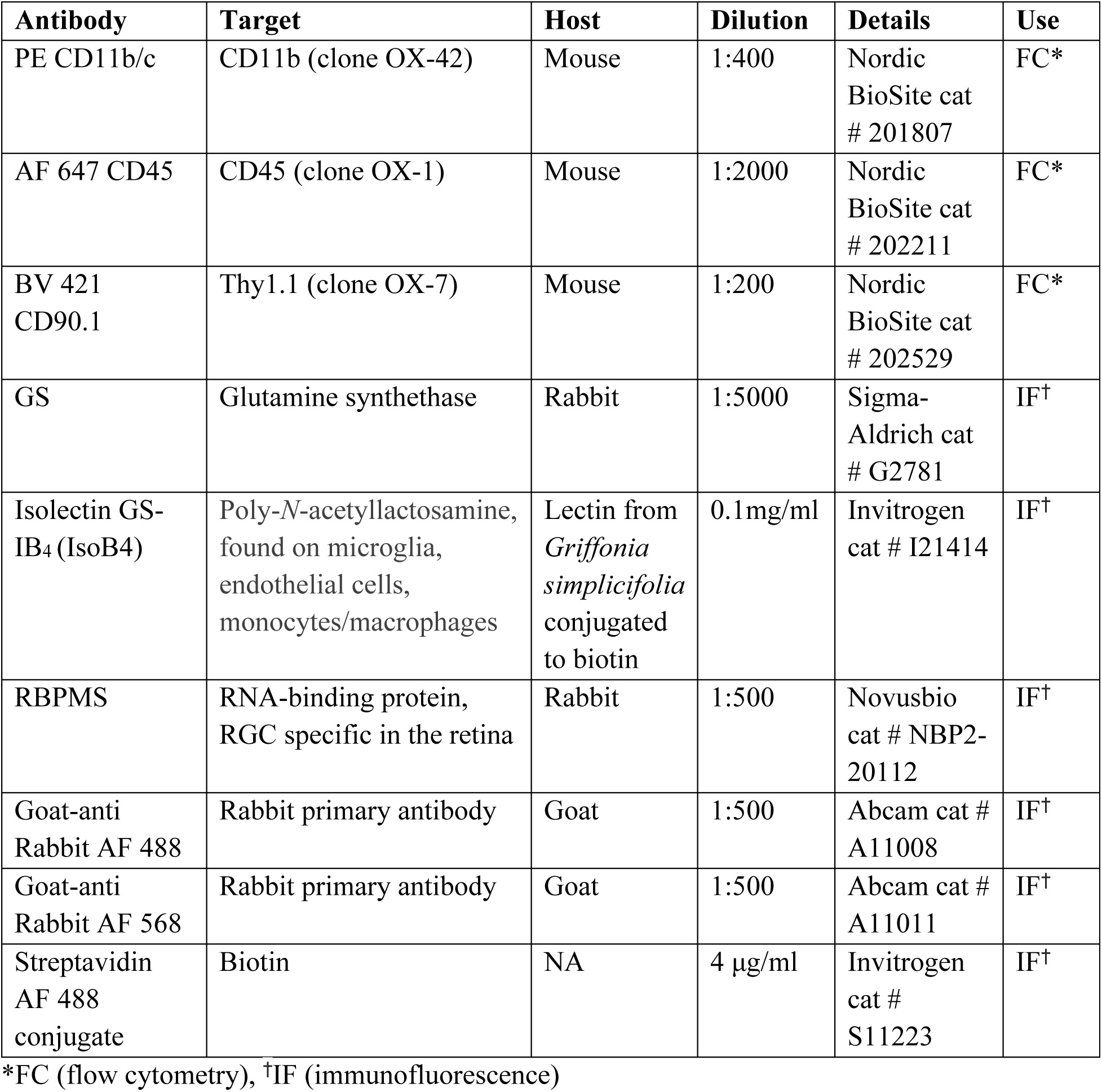
Antibody details.

### Assessment of neurodegeneration following ocular hypertension

Eyes for flat mount retina preparations (*n* = 6 NT/NT eyes, 6 OHT/OHT eyes, 5 NT/OHT eyes, 5 OHT/NT eyes all at 14 day time point were maintained in fixative as globes for 2 h before the retina was dissected free at the optic nerve head, the vitreous removed, and the retina flat mounted ganglion cell layer up on a coated glass slide (Superfrost+, Thermo Fisher). Retinas were labelled with antibodies against RBPMS and Isolectin B4, and counterstained with DAPI. Flat mounted retinas were imaged on a Zeiss Axioskop 2 plus epifluorescence microscope for quantification of retinal ganglion cell densities. Six images per retina (40X magnification, 0.25 μm/pixel) were taken equidistant to the optic nerve head (1000 μm eccentricity). Images were cropped to 150 x 150 μm and RBPMS+ cells and counted using the cell counter plugin for Fiji (24). Cell counts were averaged across the 6 images, and expressed as a density per 0.01 mm^2^.

### Microglia imaging and quantification

All microglia images were acquired on a Zeiss LSM800-Airy (20X, 319.45 x 319.45 μm, 0.312 μm/pixel, *z*-stack). Imaging parameters were kept constant to allow comparison of microglial volume. For flat mounted retinas labelled with Isolectin B4, RBPMS, and DAPI, 4 images were taken at 1500 μm superior, nasal, inferior, and temporal to the optic nerve head. In longitudinal optic nerve sections, Isolectin B4 was imaged proximal to the eye and proximal to the chiasm. Images of Isolectin B4, Nissl, and DAPI labelled brain sections were acquired at the midpoint of the SC and dLGN respectively. Microglia from retina and brain regions were reconstructed manually in Imaris (Bitplane) using the Filaments tool with automatic volume filling and *z*-depth. Parameters were kept constant for direct comparison of volumes. Microscopy, reconstructions, and image analysis were performed independently by 3 independent researchers and final reconstructions curated by an independent researcher. In the retina, microglia were classified to either the nerve fibre layer/ganglion cell layer (NFL/GCL) or inner plexiform layer based on *z*-depth, using the lower boundary of RBPMS nuclei as the limit of the ganglion cell layer. In the retina, ∼9 microglia per image were reconstructed in the NFL/GCL on average (*n* = 137 NT/NT, 185 NT/OHT, 197 OHT/NT, 222 OHT/OHT) and ∼7 microglia per image in the inner plexiform layer on average (*n* = 92 NT/NT, 134 NT/OHT, 160 OHT/NT, 170 OHT/OHT). In the brain, ∼5 (SC) and ∼6 (dLGN) microglia were reconstructed per image on average (SC, *n* = 103 NT/NT, 114 NT/OHT, 115 OHT/NT, 121 OHT/OHT; dLGN, *n* = 123 NT/NT, 116 NT/OHT, 138 OHT/NT, 130 OHT/OHT). For individual microglia, total number of branch points, total process length, total process volume, and field area were exported. Total process volume was normalized to total process length in order to distinguish microglia with large but thin processes from those with short but thick processes. Sholl analysis was performed with the soma center as the origin point and an intersection distance of 3 μm. The area under the Sholl curve (Sholl AUC) was calculated. Individual microglia were grouped by unsupervised hierarchical clustering (HC) in order to define populations of increasing reactivity. HC was performed using Morpheus (https://software.broadinstitute.org/morpheus) where every microglia from the GCL was classified based on the 5 morphological measurements described above. For HC, microglia were clustered using Euclidean distance (linkage method = average). In longitudinal optic nerve sections an area of 312(*x*) x 624(*y*) x 12.25(*z*) μm^3^ was cropped (to avoid including the optic nerve sheath). The whole channel corresponding to Isolectin B4 was reconstructed as a volume using the Surfaces automatic volume analysis tool in Imaris. Image thresholding was kept constant. Because adjacent cells were sometimes considered as one object by the Surfaces algorithm, the number of microglia within the same area was counted manually using the Imaris Spots tool.

### Quantification of infiltrating and reactive immune cells

Microglia and monocyte density was calculated by counting all microglia and monocytes in the image volume, taking the average of the 4 regions imaged, and expressing as *n*/0.001 mm^3^ (*n* = 6 NT/NT eyes, 6 OHT/OHT eyes, 5 NT/OHT eyes, 5 OHT/NT eyes; *n* = 3 brains [6 independent hemispheres] for NT/NT, OHT/OHT, NT/OHT, and OHT/NT)). Microglia in the retina were classified to the NFL/GCL or inner plexiform layer as above. Microglia and monocyte soma center positions were recorded and related to RBPMS+ nuclei centers in the retina. For each microglia, nearest neighbor distances (NND) to the nearest microglia and neuron (RBPMS+ in the retina) were calculated in R using the *nndist* function in the spatstat package (25). Monocyte NND was calculated in the same way. Gliosis was determined by glutamine synthetase (GS) staining in the retina (cryo-sections; *n* = 3 eyes for NT/NT, NT/OHT, OHT/NT, and OHT/OHT). GS labelling were quantified by average pixel intensity and space filling assessed by fractal measurement (Fractal box count).

### Flow cytometry analysis of microglia and monocyte populations

To assess retinal microglia and monocytes numbers, at day 3 rats were euthanized as above (*n* = 7 NT/NT eyes, 8 OHT/OHT eyes), whole retinas dissected free from eyes cups under ice-cold HBSS, and dissociated in dispase in 1x HBSS (Corning) at 37 °C and 350 RPM on an Eppendorf ThermoMixer C (Eppendorf). Cell were blocked with 1% BSA in 1x HBSS for 1 h and stained with antibodies against CD11b/c, CD45, and CD90.1. Cell numbers were assessed on a BD Influx equipped with 488, 561, 641 and 405 nm lasers. Gates were set based on FS *vs* SS and viability dye exclusion (LIVE/DEAD Aqua, Invitrogen). Microglia were identified as CD11b/c^+^/CD45^lo^/CD90.1^-^ cells and monocytes identified as CD11b/c^hi^/CD45^hi^/CD90.1^-^ cells. Singlets were discriminated based on FS area *vs* pulse width. Cell samples were run for 600 s while mixing (accounting for approx. 25% of total tissue homogenate) and data analyzed using FlowJo (FlowJo LLC).

### Cytokine Assay

Cytokine array analysis was performed in order to identify retinal or circulating factors that might influence immune activation in OHT and NT/OHT eyes. Rats were euthanized at day 14 and eyes enucleated (*n* = 4 eyes for NT/NT, OHT/OHT, NT/OHT, and OHT/NT). Whole retina (without optic nerve head) were dissected in HBSS and lysed in HBSS with protease inhibitors by ultrasonication (Vibra-Cell; Sonics & Materials). Samples were frozen at -80 °C overnight, thawed, centrifuged, and protein quantification performed by Bradford assay. The array (Proteome profiler rat XL cytokine array kit; R&D systems) was performed according to manufacturer’s instructions. 100 μg of protein was used for each sample and final membranes exposed to *x*-ray film for 2 s, 5 s, 10 s, 30 s, 1 min, 2 min, 5 min, 10 min, 20 min, and overnight (over-exposure). Developed films were digitized and analyzed by densitometry after background subtraction (Fiji). Spots were analyzed in duplicates and normalized to the average of the 3 reference spot duplicates. The best exposure was determined by the strength of signal and lack of spot blurring.

### Statistical analysis

All statistical analysis was performed in R. Data were tested for normality with a *Shapiro Wilk* test. Normally distributed data were analyzed by *Student’s t*-test or *ANOVA* (with *Tukey’s HSD*). Non-normally distributed data were assessed using a *Kruskal Wallis* test followed by multiple pairwise *Wilcoxon* tests with *Benjamini and Hochberg* correction. Unless otherwise stated, * = *P* < 0.05, ** = *P* < 0.01, *** *P* < 0.001, NS = non-significant (*P* > 0.05).

## Results

### Unilateral glaucoma does not cause increased intraocular pressure or retinal ganglion cell death in the contralateral eye

There was no significant difference between unilateral and bilateral normotensive eyes (NT), or between unilateral and bilateral ocular hypertensive eyes (OHT) in terms of magnitude or duration of IOP increases (**Figure 1A-B)**. OHT resulted in significant loss of RGCs as identified by RBPMS labelling. RGC density was reduced in both OHT/NT and OHT/OHT eyes compared to NT/NT and NT/OHT eyes, with no significant unilateral or bilateral effect in RGC death (**Figure 1C)**.

### Retinal microglia demonstrate increasingly reactive morphologies in unilateral normotensive, unilateral ocular hypertensive, and bilateral ocular hypertensive eyes

We performed volume reconstructions of individual microglia (*n* = 1297 microglia; **Figure 2A-D**) and assessed branching complexity, field area, and volume. Individual microglia were binned by their position within the retinal layers to either nerve fiber layer/ganglion cell layer (NFL/GCL) or IPL microglia. Sholl analysis revealed a reduction in branching density (Sholl AUC) from NT/NT for NT/OHT, OHT/NT, and OHT/NT and with increasing magnitude of reduction through these groups (**Figure 2B**). This pattern of decrease was also observed in the number of branches, total process length, and field area of microglia (**Figure 2B**). Microglial volume increased in OHT eyes compared to NT eyes but there was no significant change in volume between NT/NT and NT/OHT eyes (**Figure 2B**). The data quantitatively demonstrate morphological changes consistent with activation (retraction and increased volume (26, 27)) occurring in microglia in OHT eyes. Microglia in NT eyes contralateral to OHT eyes demonstrate early reactive morphologies (retraction). In the IPL, microglial morphology largely followed the same trend (**Figure 2C-D**). Microglia from OHT eyes demonstrated significantly decreased Sholl AUC, number of branch points, total process length, and field area (**Figure 2D**). Microglial volume was increased in OHT eyes compared to NT eyes (**Figure 2D**). The contralateral response was more muted in IPL microglia with NT/NT and NT/OHT showing no significant difference in Sholl AUC, number of branch points, and total process length but significantly reduced field area and significantly increased volume (**Figure 2D**). OHT/OHT showed significant process retraction compared to OHT/NT but no significant increase in volume (**Figure 2D**), indicating that microglia in the IPL also display activated morphologies, that follow a similar pattern in magnitude of change (OHT/OHT > OHT/NT > NT/OHT > NT/NT), but to a lesser degree than microglia in the NFL/GCL.

**Figure 2:**
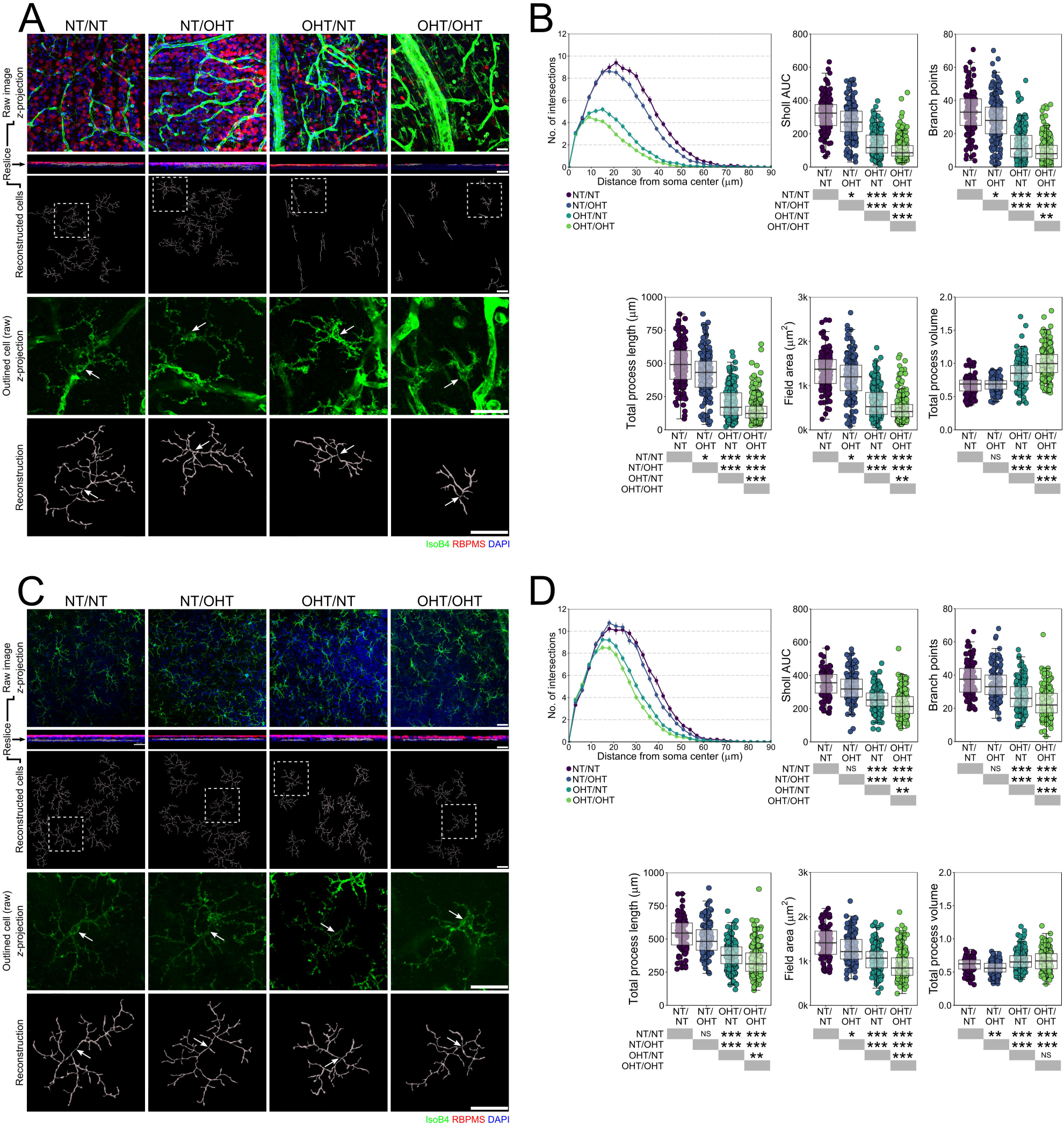
OHT results in microglia remodeling and activation. (**A**) Morphologies of individual microglia from the NFL/GCL were assessed by high resolution confocal imaging and Imaris reconstructions (*labels to left of image panel*). (**B**) Multiple metrics of microglia morphology demonstrated significant changes in microglia between all groups suggesting progressive activation from NT/NT > NT/OHT > OHT/NT > OHT/OHT (retracted processes of increased volume). (**C-D**) These data were mirrored in microglia from the IPL. For complete *n* please see **Methods**. White arrows denote microglia soma position in **A** and **C**, scale bars = 30 μm in **A** and **C**, * = *P* < 0.05, ** = *P* < 0.01, *** *P* < 0.001, NS = non-significant (*P* > 0.05).

We next performed hierarchical clustering (HC) of individual microglia in the NFL/GCL to determine whether these morphological measures alone could group microglia by degree of activation (**Figure 3**). HC produced 4 clusters: Cluster 1; a small cluster, *n* = 16, of NT/NT and NT/OHT microglia, Cluster 2; *n* = 210, predominantly consisting of NT/NT and NT/OHT microglia reflecting microglia with resting morphology (63.5% and 51.4 % of total NT/NT and NT/OHT respectively), Cluster 3; *n* = 156, a mix of microglia from all 4 groups, representing morphologies related to early activation, and Cluster 4; *n* = 359, predominantly OHT/NT and OHT/OHT microglia (67.5% and 85.6 % of total OHT/NT and OHT/OHT respectively), representing retracted and condensed morphologies consistent with activation (**Figure 3**).

**Figure 3:**
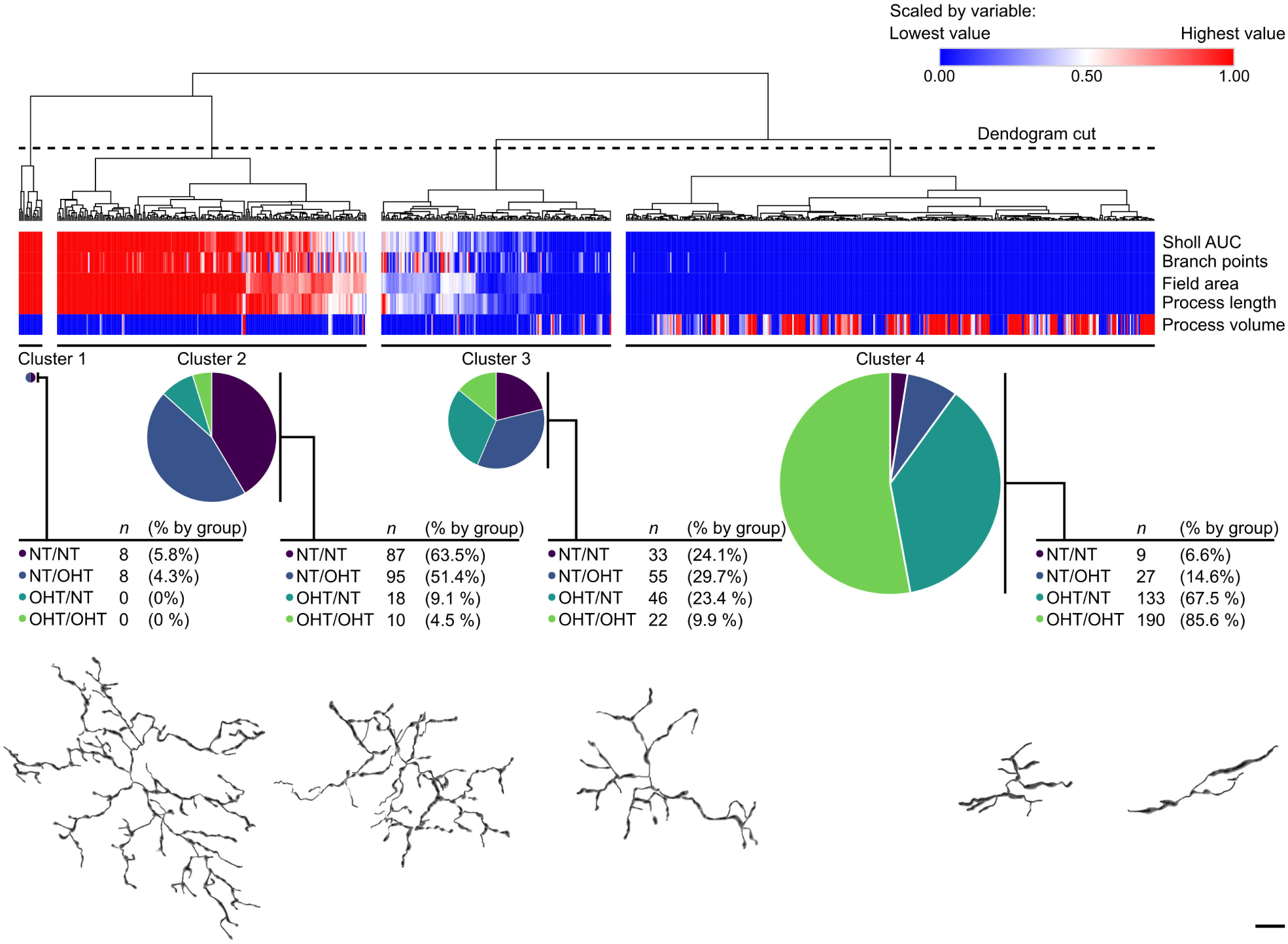
Hierarchical clustering distinguishes activation state and disease groupings. Hierarchical clustering of microglia metrics by Euclidean distance was used to cluster microglia morphologies, generating 4 independent clusters of microglia (Cluster 1-4). The breakdown of each cluster is shown *below* the heatmap/dendrogram. Example cells belonging to each cluster as shown *below* cluster breakdown. Clusters represented groups of activation and demonstrates that microglial morphological measurements can distinguish biologically meaningful groups of microglia independently of disease grouping. *Scale* bar = 10 μm.

### Monocyte infiltration occurs early following ocular hypertension

Increased numbers of microglia in the retina have been reported following OHT, as have mild increases in microglia number in NT eyes contralateral to OHT eyes. Microglia density was increased in the NFL/GCL in both OHT/NT and OHT/OHT eyes compared to NT/NT (**Figure 4A**). There was no significantly detectable contralateral effect for either unilateral NT or OHT eyes in comparison to bilateral NT or OHT eyes (**Figure 4A**). Microglia density in the IPL was less variable between groups, with only OHT/NT eyes showing a significant increase over other groups (**Figure 4A**). NND analysis revealed that the distance between microglia in the NFL/GCL was smaller on average in OHT eyes than NT eyes, with no change in NT/OHT eyes compared to NT/NT eyes (**Figure 4B**). In the IPL, NND was also smaller between microglia in OHT eyes and in NT/OHT eyes compared to NT/NT eyes (**Figure 4B**). However, the interquartile range indicated that in the IPL the majority of NNDs for OHT eyes remained overlapping with those of NT eyes, whereas in the NFL/GCL these ranges were non-overlapping (**Figure 4B**). These data suggest that changes are modest in the IPL and that microglia here are largely distributed normally, but changes are more pronounced in the NFL/GCL. Given that microglial density was substantially increased (2.8 to 3.5-fold increase), NND was not increased to the same degree (1.6 to 2.4-fold increase) demonstrating that microglia remain well distributed (*i.e.* avoid clumping/clustering). The NND between microglia and RGCs significantly increased in microglia from both the NFL/GCL and IPL under OHT in comparison to NT (**Figure 4B**). Since the increase in microglial density was concurrent with the decrease in RGC density, these data suggest that microglia are not clustering around surviving RGCs.

**Figure 4:**
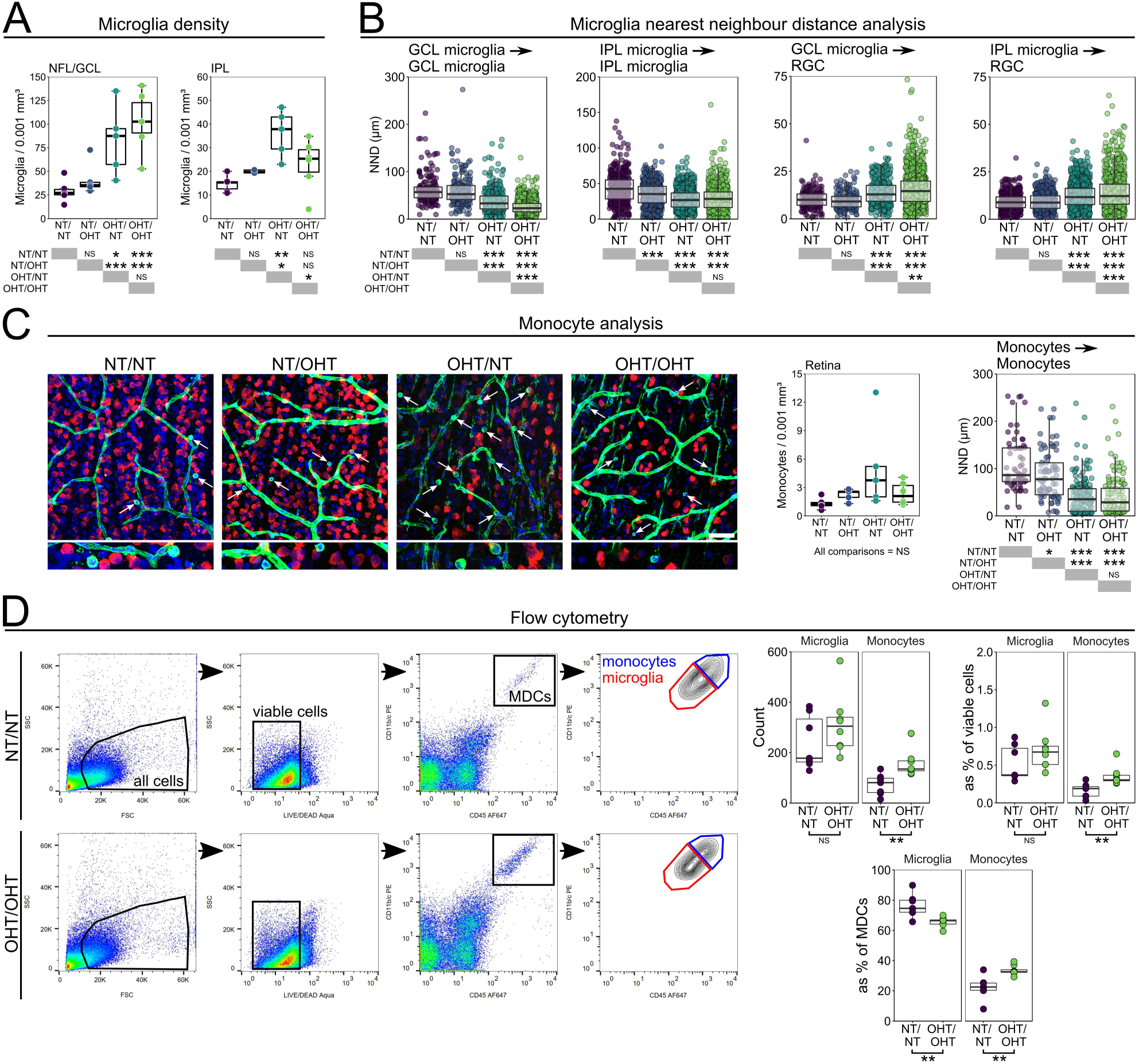
Monocytes enter OHT retina early following IOP elevation. (**A**) Following 14 days of OHT microglia numbers were increased in both the NFL/GCL and IPL of the retina. (**B**) NND analysis demonstrates that microglia get closer to each other but are diffuse around RGCs suggesting that microglia do not cluster around surviving RGCs. (**C**) Ameboid monocytes were present in all conditions after 14 days of OHT (*white arrows*) with small NNDs suggestion monocyte clumping in local regions of damage. (**D**) To begin to assess whether this increase in microglia number was due to infiltrating monocytes entering the tissue and become microglia-like monocyte derived macrophages, whole retina homogenate was assessed by flow cytometry. Retinas were sampled at 3 days post-injection (the point of peak IOP in this model). At this time point microglia numbers were not increased but there was a significant increase in monocyte numbers (both by raw count and as a percentage of other cell types). MDCs = myeloid derived cells. For complete *n* please see **Methods**. Scale bars = 100 μm in **C**, * = *P* < 0.05, ** = *P* < 0.01, *** *P* < 0.001, NS = non-significant (*P* > 0.05).

Extravasation and increased monocyte infiltration has been demonstrated as an early and persistent feature of glaucoma in animal models (28) and in post-mortem human tissue (2), and CD45^hi^ monocytes are present in the retina in low numbers under normal physiological conditions (29, 30). We observed Isolectin B4+ amoeboid cells in the majority of retinas in all experimental groups but no significant change in average monocyte density across experimental groups (**Figure 4C**). NND analysis demonstrated a significant decrease in distance between monocytes in OHT eyes compared to NT eyes (all *P* < 0.001) and a substantial shift in the interquartile range, suggesting a clumping of monocytes that may reflect recruitment rather than random transient migration into the retina (**Figure 4C**). Monocyte infiltration has been shown to be an early event in glaucoma (28). We counted retinal myeloid-derived cells (MDCs) by flow cytometry at an early time-point (3 days post-OHT induction, representing peak IOP in the absence of RGC death in our experimental groups; **Figure 4D**). Monocytes, but not microglia, were significantly increased at this early time-point in OHT compared to NT (*P* < 0.01; **Figure 4D**). This demonstrates early monocyte infiltration in the absence of microglial proliferation. This early increase in monocytes and a late increase in microglia supports a hypothesis in which infiltrating monocytes become monocyte derived microglia-like macrophages during glaucoma pathogenesis.

### No detectable Müller glia activation following ocular hypertensive glaucoma

Müller glia provide trophic and metabolic support to RGCs during times of stress, including glaucoma related stresses (31, 32). Müller glia activation, identified by increased glutamine synthetase (GS) labelling, has been demonstrated in different animal models of glaucoma. To determine whether Müller glia activation was a component of glaucomatous neurodegeneration in this model, cryo-sections were labelled with antibodies targeting GS (**Figure 5A**). Analysis of GS labelling in the NFL/GCL/IPL (representing Müller glia association to RGC axon, soma, and dendrites) did not indicate significant Müller glial activation at this time point (**Figure 5B**).

**Figure 5:**
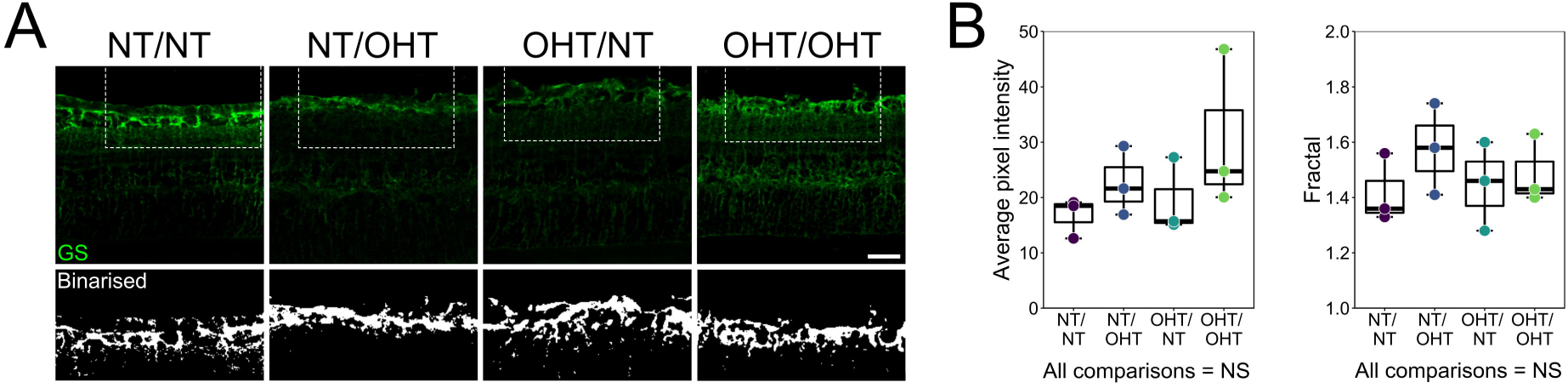
No activation of Müller glia identified by glutamine synthetase labelling. (**A**) Müller glia were labelled in cryo-sections by glutamine synthetase (GS). GS content was assessed in the NFL/GCL/IPL by measuring average pixel intensity and by fractal analysis of binarized crops. (**B**) No significant difference in average pixel intensity or fractal was observed across groups, suggesting no gross Müller glia activation at this time point in this model. Scale bars = 50 μm in **A**, NS = non-significant (*P* > 0.05).

### Microglial activation extends into the optic nerve

Since RGC axonal loss with a concomitant loss of axon transport extended into the optic nerve in this model, we sought to determine whether this was accompanied by microglial activation. Microglia were assessed in longitudinal optic nerve sections (**Figure 6A**). Microglial counts demonstrated a significant increase in microglial density both proximal to the eye and proximal to the chiasm in OHT optic nerves compared to NT optic nerves (**Figure 6B**). No contralateral change was observed in microglial density in optic nerves (**Figure 6B**). Volume reconstructions of whole Isolectin B4 content demonstrated the same trend, with significant increases in OHT optic nerves compared to NT nerves, which was more pronounced proximal to the chiasm (**Figure 6B**). Microglial activation was most evidently demonstrated at the chiasm where RGC axons from unilateral OHT eyes decussate, with the appearance of activated microglia following both optic tracts post-chiasm (**Figure 6C**).

**Figure 6:**
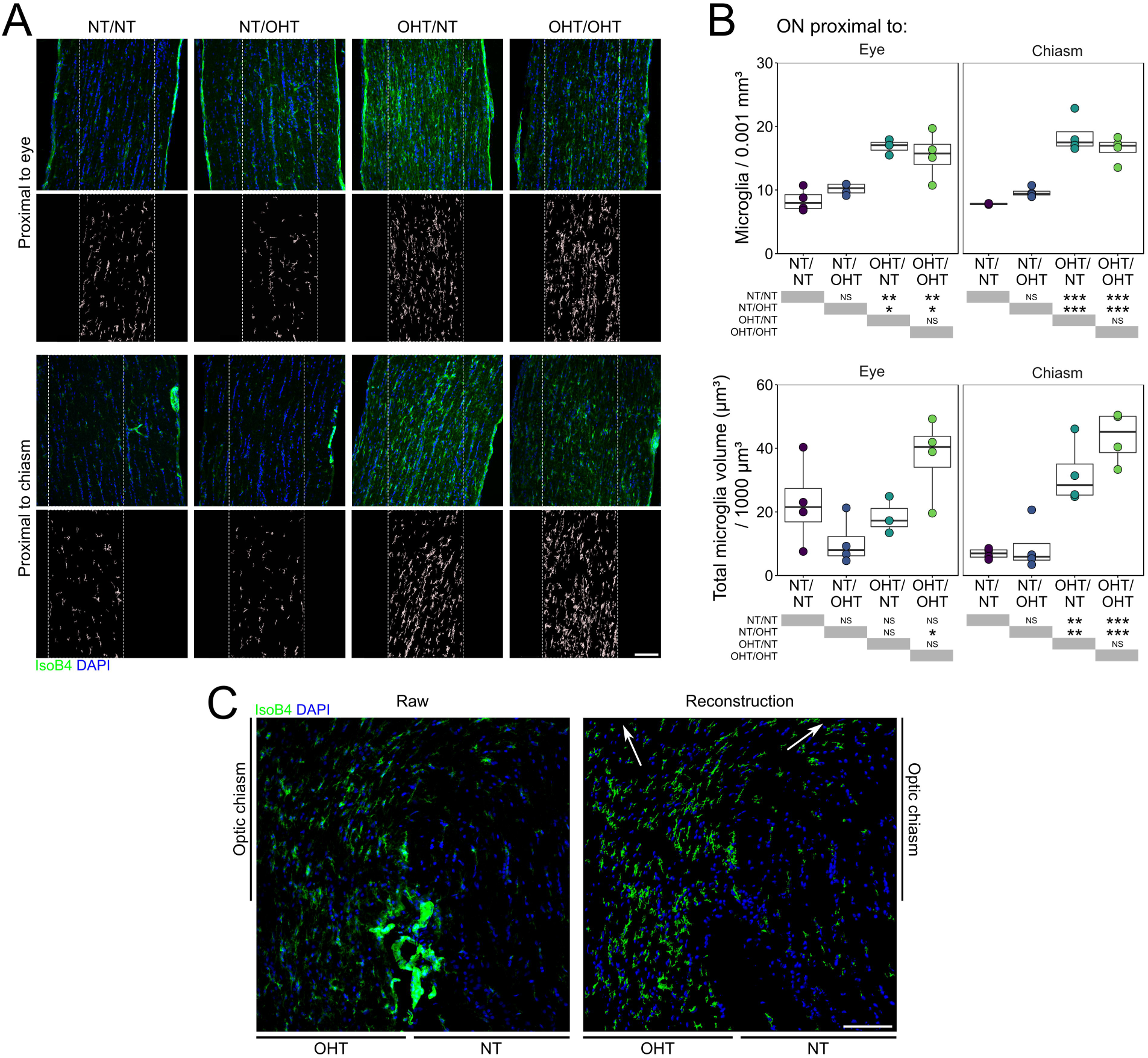
Microglia activation extends into the optic nerve and optic chiasm. (**A**) Optic nerves were transversally sectioned and overall microglial labelling was reconstructed (*insets*) and analysed. (**B**) There exists significant microglia activation in the optic nerve proximal to the eye and proximal to the chiasm. (**C**) Microglial activation is most evidently demonstrated at the chiasm where RGC axons from unilateral OHT animals decussate, with the appearance of activated microglia following both optic tracts post-chiasm (*white arrows* in reconstruction; *right*). Scale bars = 100 μm in **A** and **C**, * = *P* < 0.05, ** = *P* < 0.01, *** *P* < 0.001, NS = non-significant (*P* > 0.05).

### Microglia are activated in the dorsal lateral geniculate nucleus and superior colliculus in both hemispheres, irrespective of unilateral or bilateral ocular hypertension

Microglial activation in retinothalamic projections has been reported in animal models of glaucoma (20, 21). Brains were serial sectioned and microglia in central dLGN and SC were labelled (**Figure 7A-C**). Microglial activation was evident in the dLGN across both hemispheres in all conditions compared to brains from NT/NT animals. Microglia demonstrated reduced branching complexity, number of branch points, total process length, and field area, and an increased volume (**Figure 7D**). The dLGN contralateral to NT/OHT eyes (majority RGC input from NT eye) demonstrated significant changes in these measures compared to NT/NT dLGN, but not to its fellow dLGN (contralateral to OHT/NT eyes; **Figure 7D**), indicating that inflammation was not restricted, nor proportional, to the degree of RGC decussation. Microglia counts demonstrated no significant increase in microglia density in the dLGN, suggesting activation in the absence of proliferation or infiltration (**Figure 7D**). In the SC, microglial morphology was also indicative of activation, with microglia demonstrating a significant reduction in branching complexity, number of branch points, total process length, and field area, and an increased volume in both hemispheres from all conditions compared to NT/NT (**Figure 7E**). In the SC, there was no significant difference in these measurements in microglia between hemispheres in unilateral animals, consistent with the findings in the dLGN (**Figure 7E**). Microglia from OHT/OHT animals and both hemispheres of the unilateral animals were more similar in morphology than in the retina (**Figure 7D-E**), suggesting that the degree of microglial activation in the brain is not contingent on the degree of dysfunctional and/or absent RGC input. Microglial density was increased in the SC in OHT/NT eyes, and highly variable in OHT/OHT eyes, suggesting proliferation or recruitment of microglia in the SC (**Figure 7E**). Since amoeboid monocytes were not observed in either the SC or dLGN, and that the SC and dLGN are not directly insulted by elevated IOP, microglial proliferation is the most likely scenario. Both unilateral and bilateral OHT result in consistent microglial activation across both hemispheres of the visual thalami (*i.e.* neuroinflammatory signatures propagate even when only a small percentage of axons are damaged or dystrophic).

**Figure 7:**
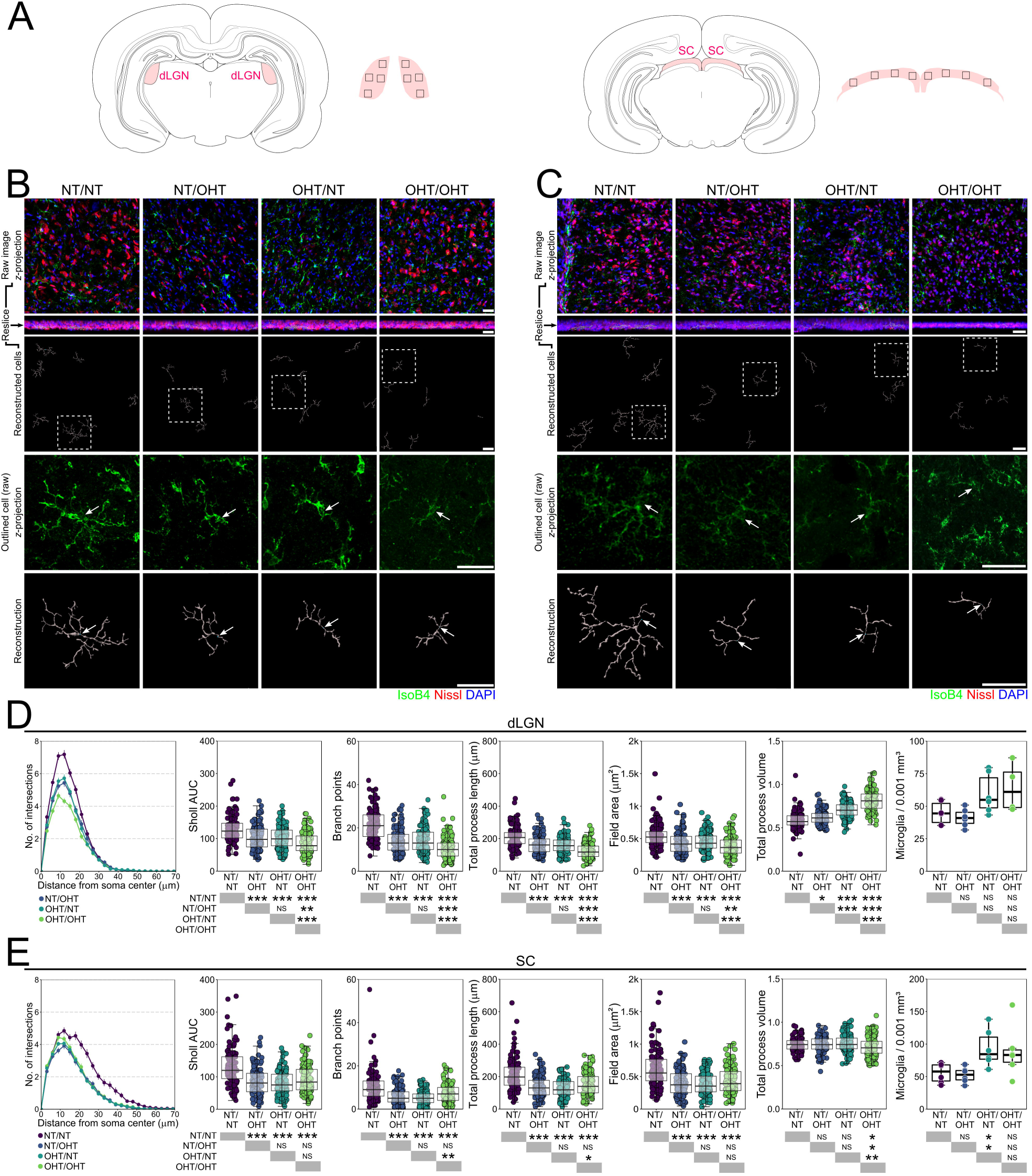
Microglia activation persists into terminal visual thalami following OHT. (**A**) Microglia were assessed from dLGN and SC sections. Regions of images are shown by black inset boxes in the schematics. Morphologies of individual microglia from the dLGN (**B**) and SC (**C**) were assessed by high resolution confocal imaging and Imaris reconstructions (*labels to left of image panel*). (**D-E**) Multiple metrics of microglia morphology demonstrated significant changes in microglia morphology between NT/NT and all other groups. These results were more binary (*i.e.* damage to a single retina drives neuroinflammatory insults to both terminating hemispheres) suggesting that neuroinflammatory signatures propagate even when only a small percentage of axons are damaged or dystrophic. For complete *n* please see **Methods**. White arrows denote microglia soma position in **B** and **C**, scale bars = 30 μm in **B** and **C**, * = *P* < 0.05, ** = *P* < 0.01, *** *P* < 0.001, NS = non-significant (*P* > 0.05).

### Upregulation of cytokines in ocular hypertension and in contralateral normotensive eyes

RGC dysfunction or neurodegeneration in the NT/OHT eyes, but with substantial microglial inflammation in the optic nerve and in terminal brain thalami, whole retinal lysate was probed for cytokine signals that could contribute to microglial activation (**Figure 8A**). Of the 79 analytes assessed by cytokine array, 6 were identified as significantly altered across groups by *MANOVA* (**Figure 8B-C**). These analytes were CNTF, Fetuin A, FGF-1, Galectin-3, ICAM-1 (CD54), and Lipocalin-2. OHT eyes demonstrated a significant upregulation of CNTF compared to NT eyes. Fetuin-A and ICAM-1 were significantly increased in OHT/NT eyes compared to NT/NT eyes (**Figure 8C**). Galectin-3 and Lipocalin-2 were significantly increased in OHT/OHT eyes compared to NT/NT eyes (**Figure 8C**). FGF-1 was significantly increased in NT/OHT eyes compared to NT/NT eyes (**Figure 8C**).

**Figure 8:**
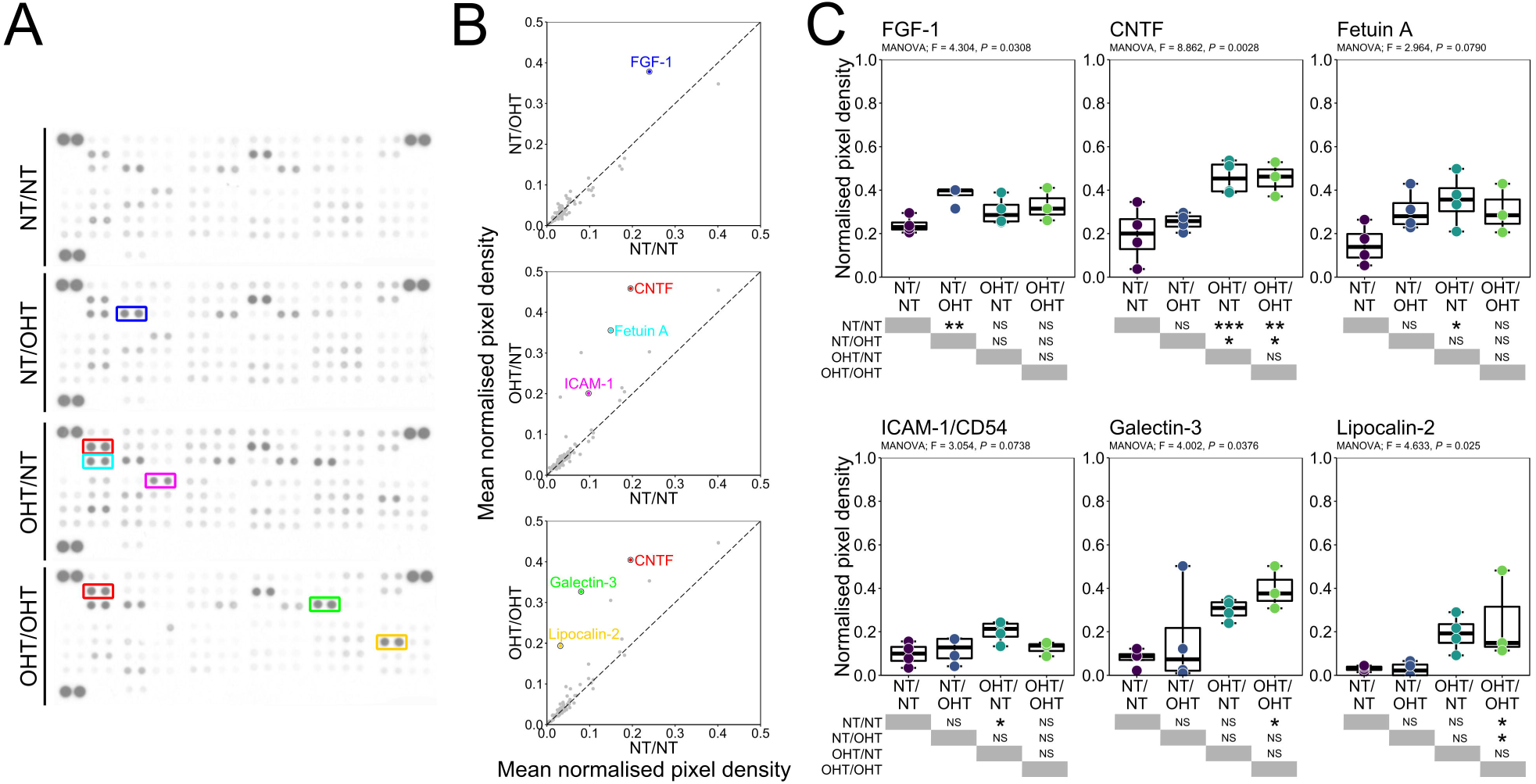
OHT drives neuroinflammatory cytokine expression in the retina. (**A**) To assess cytokine and chemokine expression in the retina following OHT, retinas were homogenized and assessed by protein array. (**B-C**) Densitometry analysis identified 1 changed cytokine (FGF-1) between NT/NT and NT/OHT groups, 3 changed cytokines (CNFT, Fetuin A, ICAM-1) between NT/NT and OHT/NT groups, and 3 changed cytokines (CNTF, Galectin-3, Lipocalin-2) between NT/NT and OHT/OHT groups. All cytokines were increased compared to NT/NT groups and may represent ideal candidates for further experimentation or as biomarker candidates. * = *P* < 0.01, ** = *P* < 0.05, *** *P* < 0.01, NS = non-significant (*P* > 0.05).

## Discussion

Retinal ganglion cells are reliant on supportive glia throughout their long projections to terminal visual thalami (33). Following periods of elevated IOP these glia can become reactive, likely as an initial protective response that actually further compromises retinal ganglion cell survival (3, 34). Supporting this, complete ablation of microglia or monocytes only provides short-term neuroprotection in glaucoma (28, 35). In a recent study, almost complete depletion of resident microglia and infiltrating monocytes (using a PLX5622 and clodronate liposome combination treatment) failed to provide benefit in a more acute model (controlled optic nerve crush) and actually delayed axon regeneration following lens injury-induced RGC axon regeneration (36). In the present study, quantification of microglial morphology demonstrates retracted morphologies and increased process volume consistent with activation from a resting state (26, 27). Activated morphology persisted throughout the retina, optic nerve, and brain mirroring retinal ganglion cell dysfunction that extended along the trajectory of the axon. Reciprocal signaling between microglia and neurons plays an important role in defining microglia phenotypes (37). The resting, non-activated state, is in part controlled by CD200 and fractalkine (CX3CL1) signaling. CD200 is expressed by neurons and through binding CD200R on microglia, maintains a resting state (38, 39). CX3CL1 is expressed in neurons and can be membrane bound and in a secreted form; it signals through its receptor on microglia to maintain a resting state (40, 41). CD200 and CD200R expression changes early in glaucoma while loss of CX3CL1 signaling causes earlier axon transport dysfunction and in a DBA/2J mouse model of glaucoma (42, 43). Neurons also express markers of injury (*e.g.* surface expression of CCL21 which activates CXCR3 on microglia (39, 44, 45)) or of health (*e.g.* surface expression of CD47, the ‘don’t eat me’ signal (46, 47)). Activated microglia exhibit increased expression of toll-like receptors, complement receptor and components, and pro-inflammatory cytokines and chemokines. Microglia and complement involvement in synapse elimination during the critical period and in neurodegenerative disease is well established (48-50). Complement components are upregulated in glaucoma in animal models and in human tissue, which play a critical involvement in synaptic pruning in glaucoma, and altered retinal ganglion cell survival through its manipulation (51-58). This reciprocal relationship between microglial activation and neuronal dysfunction may underlie why microglial activation is so robust in the SC and dLGN even in the absence of direct insult from OHT (*e.g.* direct IOP and vascular compromise).

The degree to which unilateral OHT affects the contralateral eye requires further elucidation. We confirmed quantitatively the observation of activated microglia in NT eyes contralateral to OHT eyes. In these eyes, as in the brain, activation occurs in the absence of direct OHT insult. The source of microglial insult is therefore either systemic or neuron-derived. We observed no significant change in IOP in NT/OHT eyes over NT/NT eyes. We also demonstrated no detectable retinal ganglion cell loss in the retina in NT/OHT eyes. In the rat, decussation at the chiasm of ∼90% of axons to the SC and dLGN contralateral to the eye of origin leaves ∼10% of RGC axons projecting ipsilaterally. Significant microglial activation therefore persists along a substantial portion of these retinal ganglion cell axons and presents a pro-neuroinflammatory environment to otherwise healthy retinal ganglion cell terminals. It is possible that this interaction translates into activation at the somal end in the retina, although this remains to be experimentally validated. Of note is the contralateral effect observed as the difference in activation between unilateral and bilateral OHT in the retina. This furthers the hypothesis that cumulative neuronal insult or systemic effects may drive activation to a greater degree in the retina. This graded effect in the retina was not observed in the brain, where microglial activation was consistent across hemispheres irrespective of whether the majority of input was from an OHT or NT eye in unilateral animals. The degree of microglial activation, and its implications towards neuro-glial signaling, neuroinflammatory environment, and retinal ganglion cell health should caution against the use of the contralateral eye as a control in unilateral OHT animals.

Microglial proliferation and migration and monocyte infiltration have been reported in a number of glaucoma models and in human glaucoma (2, 12, 28, 35, 43, 59-62). We demonstrate that an early increase in monocytes, but not microglia, and a late increase in microglia, but not monocytes occurs following sustained periods of OHT. These data suggest that monocytes infiltrate the retina and become microglia-like monocyte derived macrophages, as has been reported in Alzheimer’s disease and other neurodegenerative diseases and in retinal photoreceptor degeneration (63-67). Definitively testing this hypothesis would require complex reporter alleles that are currently unavailable in the rat and therefore outside the scope of this study. We have previously shown that vascular permeability is compromised in glaucoma, leading to increased extravasation of monocytes in addition to recruitment through vascular endothelial changes (28, 62).

In our model, increased microglial activation and monocyte infiltration results in a significant pro-inflammatory environment. Cytokine and chemokine profiling in the retina identified a number of changes in OHT and in contralateral NT eyes. CNTF, Fetuin A, FGF-1, Galectin-3, ICAM-1 (CD54), and Lipocalin-2 were all differentially elevated. CNTF, which can be released from microglia as a response to central nervous system injury, has demonstrated neurotrophic properties and increases retinal ganglion cell survival in glaucoma models (68, 69). CNTF also induces gliosis in Müller cells and leads to upregulation of pro-inflammatory cytokines and chemokines (70-72), and as such its action is likely highly context dependent. Fetuin-A, a serum carrier protein, has been proposed as a biomarker of systemic inflammation, vascular disease, and neurodegenerative disease (73-75). Reduced expression of Fetuin-A correlates with the severity of cognitive impairment in Alzheimer’s disease and Fetuin-A deficient mice demonstrate protection in experimental models of multiple sclerosis (76, 77). Galectin-3 mediates cell migration and adhesion, inflammatory cytokine release and apoptosis and has been shown to activate microglia, astrocytes, monocytes, and other immune cells (78). Plasma levels of Galectin-3 correlate with Huntington’s disease severity in patients and animals, and are associated with adverse outcomes in stroke and cerebral infarction (79, 80). Expression of Galectin-3 was increased in Huntington’s disease and genetic ablation of *Lgals3* is protective in animal models (79). Microglial released Galectin-3 can act as a ligand for TLR4, sustaining inflammatory responses (81). Increased expression of Galectin-3 has been detected in human glaucomatous donor tissue in the trabecular meshwork and optic nerve head in association with increased fibrosis (82). Icam-1 is a glycoprotein that is expressed on leukocytes, macrophages and vascular endothelium. Its expression is upregulated in response to inflammatory stimuli (IL-1, TNF-α) where it functions as a ligand for integrins to facilitate endothelial binding and transmigration of leukocytes (83). *ICAM1* expression is increased in glaucoma patient leukocytes but not blood plasma (84, 85). The upregulation of *Icam1* is early and persistent in the optic nerve head and retina of DBA2/J mice, and is reduced by radiation treatment which is highly protective for retinal ganglion cells (35). Lipocalin-2 is an antibacterial protein (acting through iron-sequestration) that elicits pro-inflammatory responses and is implicated in a number of inflammatory and neurodegenerative diseases, including Stargardt disease and age-related macular degeneration in the retina (86, 87). *Lcn2* expression is also highly upregulated in DBA2/J glaucoma and in the rat hypertonic saline glaucoma model (35, 88). A recent GEO screen identified *LCN2* involvement in glaucoma pathogenesis (89). Increased FGF-1 expression was identified in NT/OHT eyes. FGF-1 has demonstrated neuroprotective responses in neurodegenerative disease, but also activates astrocytes in the spinal cord, leading to pro-inflammatory responses (90-93). Whether this represents a neuroprotective or inflammatory response related to microglial activation warrants further exploration. These pro-inflammatory molecules represent ideal candidates for further exploration using targeted knockouts, gene therapies, or antibody therapies in glaucoma.

## List of abbreviations

AUC: (area under the curve),
FC: (flow cytometry),
GCL: (ganglion cell layer),
H&E: (haematoxylin and eosin),
HC: (hierarchical clustering),
IF: (immunofluorescence),
IHC: (immunohistochemistry),
IOP: (intraocular pressure),
IPL: (inner plexiform layer),
MDC: (myeloid-derived cell),
NND: (nearest neighbor distance),
NT: (normotensive),
NFL: (nerve fiber layer),
OHT: (ocular hypertension).

## Declarations

### Ethics approval

All experimental procedures were undertaken in accordance with the Association for Research for Vision and Ophthalmology Statement for the Use of Animals in Ophthalmic and Research. Individual study protocols were approved by Stockholm’s Committee for Ethical Animal Research (10389-2018).

### Consent for publication

N/A

### Availability of data and material

All data generated or analyzed during this study are included in this published article.

## Acknowledgments

The Authors would like to thank Monica Aronsson and Diana Rydholm for their assistance with animal husbandry and maintenance, St. Eriks Eye Hospital for financial support for research space, clinical histopathology, and animal facilities, the Knut and Alice Wallenberg Foundation and Karolinska Institutet for supporting the CLICK imaging facility, Annika van Vollenhoven at the Flow Cytometry Core Facility at the Karolinska Institutet Center for Molecular Medicine for flow cytometry, Charlotte Taul Brændstrup for assistance with Müller cell immunofluorescence, B Paul Morgan for critical reading and editing of the manuscript, and CITER (Cardiff Institute of Tissue Engineering & Repair) for supporting EK in the form of a research travel award.

## Author contributions

*JRT* – designed and performed experiments, analyzed data, wrote the manuscript; *EK* – designed and performed experiments, analyzed data, wrote the manuscript; *AO* – performed experiments, analyzed data; *FP* – performed experiments, analyzed data; *EL* - performed experiments; *RV* – performed experiments; *MK* - provided resources, wrote the manuscript; *HA* - provided resources, designed experiments, wrote the manuscript; *JEM* – provided resources, designed experiments, wrote the manuscript; *PAW* – conceived, designed, performed experiments, analyzed data, wrote the manuscript. All authors read and approved the final manuscript.

## Commercial relationships disclosures

The Authors report no commercial relationships or competing interests.

